# An optimized appetitive visual short-term memory paradigm in *Drosophila*

**DOI:** 10.1101/2025.02.28.640815

**Authors:** Brandon L. Holder, Jennifer A. McEllin, Stephane Dissel

## Abstract

The ability to generate and recall memory is a behavior that is evolutionarily conserved across the animal kingdom from humans to jellyfish. Memory not only allows previous experiences to inform future decision making, but it also amasses information essential to life, such as memory of quality food sources, shelter, and predator-related associations. Associative memory forms a relationship between two or more distinct and initially unrelated stimuli and can be defined by its temporal characteristics, such as short- and long-term duration, as well as the memory being appetitive or aversive, generating approach or avoidance behavior, respectively. Since its introduction as a memory model in the 1970s, the fruit fly, *Drosophila melanogaster*, has emerged as a powerful tool for the investigation of memory-related processes. While a variety of memory paradigms have been used extensively in *Drosophila*, such as appetitive and aversive olfactory memory, the use of appetitive visual memory remains infrequent. A previous study introduced a visual short-term memory (STM) paradigm that could be used for the study of both appetitive and aversive visual memory in *Drosophila*. However, this protocol required 50+ flies per condition, with three conditions per experiment, and 15 or more replications were frequently used to assess memory. As a result, this paradigm requires substantial numbers of flies, time, and is impractical for large genetic screens. Here, building upon this previous work, we describe an optimized appetite visual STM paradigm in freely moving *Drosophila*. Using recently published data on sexual dimorphism, innate color preferences, and borrowing practices from related appetitive assays, we have established an approach that minimizes confounding factors, such as sexually dimorphic starvation survival and sucrose preference, as well as pre-training color preference variation between groups. In doing so, we present an appetitive visual STM paradigm requiring substantially fewer replicates and numbers of flies to produce significant learning.

## Introduction

The ability to generate and recall memories is essential for living organisms. This is evidenced by its evolutionary conservation across the animal kingdom, from primates to zebra fish, fruit flies, nematodes, and even jellyfish (Beran et al., 2016, Reemst et al., 2023, Davis, 2023, Rahmani and Chew, 2021, Bielecki et al., 2023). Memory can be defined temporally based on its duration, including short-term memory (STM), lasting seconds to minutes and long-term memory (LTM), lasting for days and even years (Atkinson and Shiffrin, 1968, Cowan, 2008, Guskjolen and Cembrowski, 2023). It can also be classified as appetitive, in which the associative memory generates approach/attraction to a cue, or aversive, in which the cue generates avoidance behavior. The seminal work of Pavlov demonstrated that animals display associative memory via classical conditioning, where a conditioned stimulus (feeding) was linked to an unconditioned stimulus (ringing bell) (Pavlov and Anrep, 1927). As a result, the ringing of the bell generated an anticipatory feeding response in dogs, demonstrating associative memory.

Since the 1970s, the fruit fly, *Drosophila melanogaster*, has emerged as a powerful model for the investigation of learning and memory (Quinn et al., 1974, Tempel et al., 1983, Spatz et al., 1974). In 1974, the lab of Seymour Benzer published the first-ever evidence of learning and memory in *Drosophila* (Quinn et al., 1974). In this seminal work on aversive memory, two discernable odors were presented to flies, one of which was paired to an electrical shock. Flies that experienced an electrical shock paired to a specific odor would avoid that odor during subsequent periods, even in the absence of electrical shock. Around this same time, an early visual memory assay in *Drosophila* was presented, which used different wavelengths (colors) of light paired with an electric shock to generate aversive memory (Spatz et al., 1974). Nearly 10 years later, the first published appetitive memory assay demonstrated that fruit flies could form an appetitive association between an odor (conditioned stimulus) and a sucrose reward (unconditioned stimulus) (Tempel et al., 1983). This study also demonstrated that this memory formation was robust, generating LTM when tested 24 hours after training.

A later study by Schnaitmann *et al*. combined appetitive and visual memory in which flies associated a color stimulus with a sucrose reward in a STM paradigm (Schnaitmann et al., 2010). In this study, Canton-S (CS) flies were starved in mixed-sex populations prior to training to induce motivation to find food (the sucrose reward). Flies were then divided into three groups: green trained (sucrose paired with the color green), blue trained (sucrose paired with the color blue), and untrained (sucrose absent during entirety of training). Using a single post-training video to assess color preference between green and blue, significant STM was observed after 15 to 20 experiments consisting of 150 to 300 flies per experiment (50 to 100 flies per condition). However, this requires a substantial number of flies to assess STM. A later publication using this original protocol exhibited similar appetitive learning but still required 10 or more tests (Vogt et al., 2014).

While the original publication presented a well-designed and approachable appetitive visual STM protocol, upon implementation, we recognized that optimizations could be introduced to demonstrate learning with fewer tests, significantly reducing the number of flies required. Using data on sexual dimorphism, innate color preference, and practices from related appetitive assays, we developed an optimized approach to assess appetitive visual memory in freely moving *Drosophila*, requiring fewer trials and numbers of flies to produce significant learning.

## Materials and Methods

### Drosophila stocks and rearing

Flies were cultured at 25°C with 50% humidity under a 12h light:12h dark cycle. Flies were kept on a standard cornmeal diet (per 1L: 25 g inactive yeast, 15 g corn syrup, 25 g cornmeal, 66.67 g molasses, and 7 g agar).

### Starvation Survival

Male and female CS flies were collected upon eclosion and maintained in vials in same-sex only or mixed-sex (1:1 ratio) groups of 30 flies. Flies were aged for 5 days and then transferred into vials with starvation medium (1% agar) in same-sex or mixed-sex populations. Vials were checked approximately every 12 hours and the number of flies remaining alive was recorded. This process was repeated for 72 hours and the percentage of flies alive at each time point was calculated using % alive = (# of flies alive / starting # of flies) * 100.

### Female Mating Status

CS virgin females were collected and mixed with naïve but mature (>3 day old) don juan-GFP males in a 1:1 ratio, 30 total flies per vial. Random vials were selected for analysis every 24-hour period. Flies were ice-anesthetized and the number of females containing GFP-positive signals in the abdomen were counted under a fluorescent microscope. This process was repeated over 5 days, with day 5 representing the female population on the day of training and testing in the memory assay. The percentage of mated females was calculated using % mated = (# of mated females / starting # of virgin females) * 100.

### Simulated Training Video

Male and female CS flies were collected upon eclosion and kept in vials in same-sex only or mixed-sex (1:1 ratio) groups. Flies were aged for 5 days and then transferred to starvation medium (1% agar). On the day of the test, flies were ice-anesthetized and males were collected at 21 hours of starvation, while females were placed back onto starvation medium until 28 hours of starvation. Flies were then subjected to a simulated training battery where a color was paired with sucrose-infused filter paper, 1 minute on with 1 minute darkness and repeated 5 times. A computer monitor (Dell P2219H) with the brightness set to 100 was laid flat during the various color tests with PowerPoint slides in presentation mode that corresponded to each color. Flies were recorded for each of the 1-minute rounds of simulated training and the percentage of flies present on the filter paper in the presence of each color was recorded. This process was repeated for all colors (R:G:B) tested: red (200:0:0), orange (255:192:0), yellow (255:255:0), green (0:255:0), blue (0:112:192), and violet (155:38:182). The percentage of flies present on the filter paper for each color was calculated as % on filter paper = (# of flies on filter paper / total # of flies in chamber) *100.

### Starting Color Preference

Male and female flies were collected upon eclosion and kept in vials in same-sex only or mixed-sex (1:1 ratio) groups. Flies were aged for 5 days and then starved on starvation medium (1% agar). On the day of the test, flies were ice-anesthetized and males were collected at 21 hours of starvation, while females were placed back onto starvation medium until 28 hours of starvation. These groups were then subjected to the alternating green-violet color quadrant and testing (blank) filter paper and recorded for 90 seconds. PIG values were calculated using PIG = (# on green side - # on violet side) / total # of flies.

### Short-term Memory Apparatus

The apparatus for the appetitive visual short-term memory assay is adapted from the established protocol by Schnaitmann et al. (Schnaitmann et al., 2010). *Drosophila* egg-laying cages (FlyStuff 100 mm, Genesee Scientific) were enclosed in flat black foil (Rosco Cinefoil), the insides coated with 2871B Insect-a-Slip (BioQuip Products), and the steel cloth at the end was removed to create chambers for each condition. 100 mm petri dish lids were used as caps for each end of the chamber, one of which was permanently taped and used as the viewing and video recording end of the tube. On the opposite end, where filter paper was alternated during training and testing, the petri dish lids were taped along the raised edges to ensure a sufficient fit with the courtship tube. For training, 90 mm filter paper (P4, Fisher Scientific) was infused with either 1.5 mL of respective sucrose concentration or 1.5 mL of water and dried prior to training. To generate the training and testing color apparatus, Microsoft PowerPoint was used to generate training and testing slides (Supplementary Figure 1). The testing slide contained three circles, one for each training condition, divided into alternating quadrants of green and violet. The background surrounding the circles and engulfing the remainder of the slide was red. R:G:B for each color is: green (0:255:0), violet (155:38:182), and red background (200:0:0). The training slide was split in half with green on one side and violet on the other. A computer monitor (Dell P2219H) with the brightness set to 100 was laid flat during the training and testing periods with the PowerPoint slides in presentation mode.

### Video Capture and Analysis

Both pre- and post-training videos are captured at 1 fps for 90 seconds (videos are usually longer to accommodate a full 90 seconds for last chamber placed onto the testing array). Videos were captured in OBS Studio with Ausdom AW335 web cameras mounted to clamps that were placed above the top of each appetitive visual chamber. Videos were obtained in a dark room where the only light source was from the monitor that had the testing array. MKV videos were converted to MP4 using OBS Studio’s remux tool.

Appetitive visual memory videos were imported into ImageJ using the FFMPEG tool as a non-virtual stack and saved as TIFFs. Since the videos were recorded at 1 fps, each slice of the TIFF stack represents 1 second of time in which we can quantify the number of flies on each color per second over the 90 second duration. Two ImageJ macros were developed to expedite this counting, one for the green quadrants and one for the violet quadrants (see Supplementary Material for files). The polygon selection tool was used to outline the green quadrants, the green-specific macro was initiated, and the resulting count was saved as a .csv file. The violet quadrants were then analyzed using the violet-specific macro and saved as a .csv file. This process was repeated for the Green+, Violet+ and untrained conditions on both the pre- and post-training videos. A Microsoft Excel template was created to record the first frame, remove dead flies, and calculate PIG, change in PIG (ΔPIG), and learning index (LI).

### Appetitive Visual Short-term Memory Protocol

The protocol for the appetitive visual short-term memory assay is adapted from the established protocol by Schnaitmann et al. (Schnaitmann et al., 2010). Flies were collected upon eclosion and placed into mixed-sex groups with a 1:1 ratio of males to females. Flies were aged approximately 4 days and then transferred into starvation vials (1% agar and water) at 12:00pm on the day prior to training and testing. The following morning, flies were ice anesthetized and separated into same-sex groups in new starvation vials. After allowing 30+ minutes to elapse after flies were removed from ice, male flies were immediately trained for appetitive visual memory, while females were left to starve for slightly longer to increase motivation. Prior to training, flies were divided into three groups, green trained (Green+), violet trained (Violet+), and untrained and placed into the chambers described in the apparatus section directly above. A pre-training video was recorded in which each chamber is placed onto a testing circle with alternating green and violet quadrants and a piece of blank filter paper. The flies were then subjected to the training protocol, in which Green+ flies would be exposed to sucrose-infused filter paper when on green light and water-infused filter paper when on violet light. Reciprocally, the Violet+ flies would be exposed to sucrose-infused filter paper when on violet light and water infused filter paper when on green light. The untrained group would be exposed to water-infused filter paper on both green and violet light. After alternating 1-minute periods of 5 rounds of sucrose and 4 rounds of water training (starting and ending with a sucrose period), flies are transferred to a blank piece of filter paper and kept in total darkness for 5 minutes. After 5 minutes, flies are placed on the testing array to obtain a post-training video. A pre-training PIG is calculated for each condition (Green+, Violet+, and untrained) using PIG = (# of Flies on Green – # of Flies on Violet) / Total # of Flies. A post-training PIG is calculated using the same equation. The ΔPIG is calculated as ΔPIG = PIG_Post-training_ – PIG_Pre-training_. Finally, a learning index (LI) can be calculated using LI = (ΔPIG of Green+ – ΔPIG of Violet+) / 2. For the untrained group, LI = (ΔPIG of untrained – 0) / 2 is used to calculate the LI, since the expected ΔPIG within this group should theoretically average to zero.

### Statistical Analysis

Statistical analyses were performed with GraphPad Prism10 software. All data was tested for normality using the D’Agostino-Pearson test. Data that was normally distributed was analyzed with parametric statistical tests, including one-sample t tests, unpaired t tests, and one-way ANOVA tests with Dunnett’s multiple comparisons. For data that was not normally distributed, nonparametric statistics were used, including Wilcoxon signed-rank tests, Mann-Whitney U tests, and Kruskal-Wallis tests with Dunn’s multiple comparisons. For bar charts used, bars represent that mean of the data and error bars represent the SEM of the data. For box and whisker plots, the bottom of the box represents the first quartile while the top of the box represents the third quartile. The middle line within the box represents the median of the data, while the whiskers denote the 10th to 90th percentiles for the data. On all graphs, statistically significant findings are reported as ns = not significant, * = p < 0.05, ** = p < 0.01, *** = p < 0.001, and **** = p < 0.0001.

## Results

### Controlling for Sexual Dimorphism in Starvation Survival

The original protocol for appetitive visual STM trained and tested flies using mixed-sex populations without noting the ratio of males to females. The foundation of appetitive assays is the motivation to learn, which is produced by the starvation of flies, which motivates flies to locate food sources, such as sucrose (Quinn et al., 1974, Tempel et al., 1983). However, it has been shown that male and female *Drosophila* exhibit dissimilar responses to starvation, with mated males showing significantly higher mortality over time compared to females (Chauhan et al., 2021, Service, 1989). Mated males are also more susceptible to starvation when compared to naïve males, while mated females are more resistant to starvation than virgin females (Service, 1989). Confoundingly, virgin females have been shown to be more susceptible to starvation when compared to naïve males (Service, 1989, Huey et al., 2004). Since a mixed-sex population was utilized in the original assay, the starvation level, and therefore motivation level, can vary significantly both within and between groups of flies based on the age and ratio of male to female flies.

To explore how starvation survival varies within the context of the appetitive visual STM assay, we performed a starvation survival assay on Canton-S (CS) flies. We investigated three different conditions with variations in maturation and starvation states. For the first condition, flies were collected as naïve males or virgin females and underwent both maturation and starvation in same-sex groups. For the second condition, naïve male and female flies were collected and underwent both maturation and starvation in mixed-sex groups with a 1:1 male-to-female ratio. For the third condition, naïve male and female flies were collected and underwent mixed-sex maturation in a 1:1 ratio of males-to-females, followed by separation into same-sex groups for starvation. Flies were reared in respective groups for 3 days and then were transferred to vials containing a 1% agar starvation medium for the starvation survival assay.

CS male and female flies show comparable survival in same-sex maturation and starvation groups, with modest but significantly higher survival observed in females starting at 24 hours (Figure 1A). These data appear to conflict with previously published results (Service, 1989, Huey et al., 2004), however, this may be attributed to different food recipes being used between labs and the diets’ subsequently-induced differences in starvation survival, even within flies of the same genotype (Chippindale et al., 1996, Chippindale et al., 1998). Moderate differences were observed in the mixed-sex maturation and starvation groups, with females showing significantly greater survival starting at the 12-hour mark and consistently outperforming male flies (Figure 1B). The mixed-sex maturation and same-sex starvation groups exhibited the greatest dissimilarity between males and females, with males quickly deteriorating after the 12-hour mark (Figure 1C). Interestingly, female flies in this condition showed robust starvation survival.

**Figure 1:**
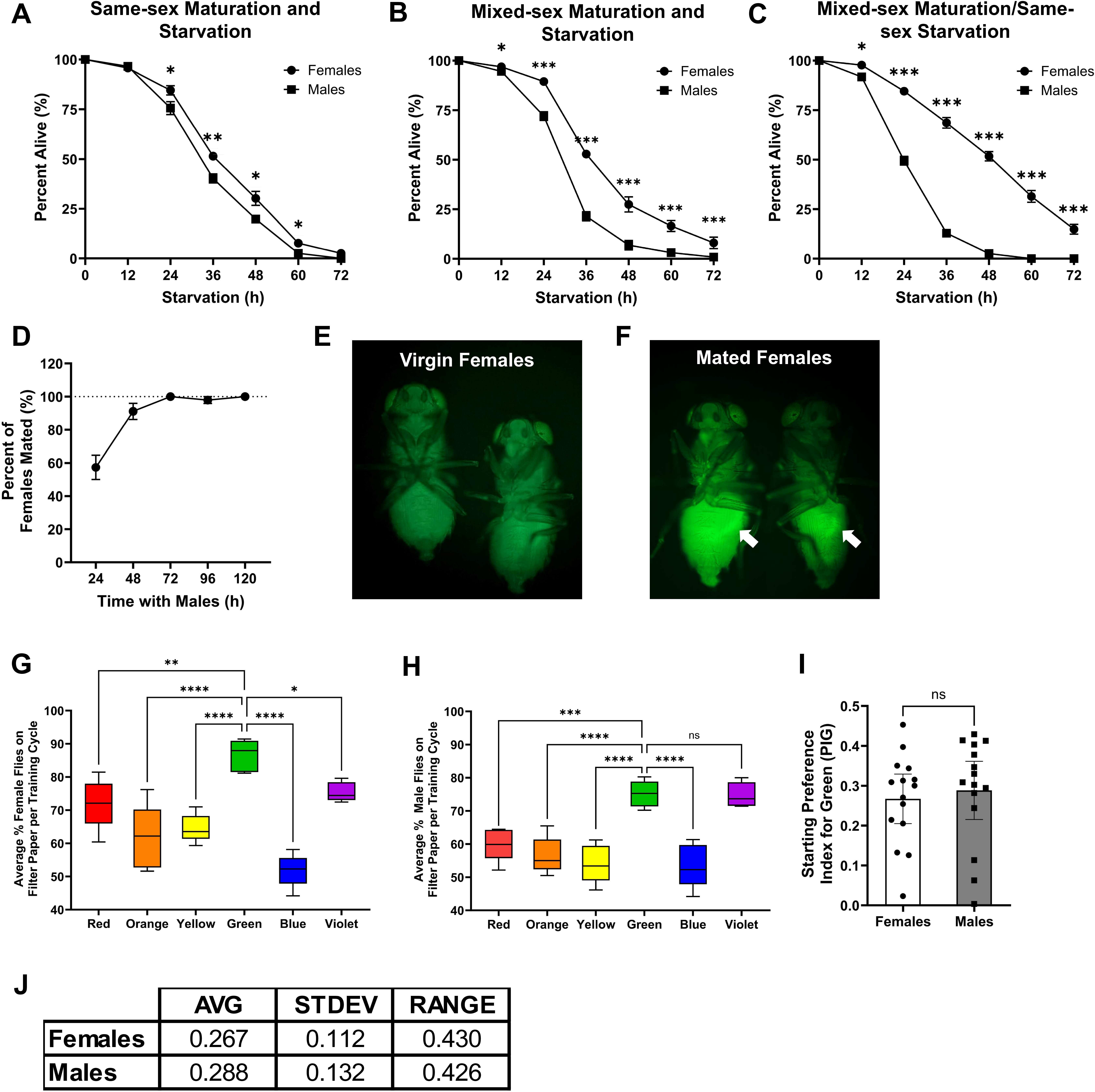
Investigating potential points of optimization for appetitive visual STM. **(A)** Starvation survival curve for CS male and female flies collected upon eclosion, reared, and starved in same-sex groupings. n = 7 groups of 50 flies per time point. Mann-Whitney test of male vs. female survival for each time point. n = 7 groups of 50 flies per time point. * = p < 0.05, and ** = p < 0.01. **(B)** Starvation survival curve for CS male and female flies collected upon eclosion, reared, and starved in mixed-sex groupings of a 1:1 male to female ratio. Mann-Whitney test of male vs. female survival for each time point. n = 7 groups of 50 flies per time point. * = p < 0.05, and *** = p < 0.001. **(C)** Starvation survival curve for CS male and female flies collected upon eclosion and reared in mixed-sex groupings of a 1:1 male to female ratio and then starved in same-sex groupings. Non-parametric t test of male vs. female survival for each time point. Mann-Whitney test of male vs. female survival for each time point. n = 7 groups of 50 flies per time point. * = p < 0.05, and *** = p < 0.001. **(D)** Percent of mated CS females as indicated by GFP-positive fluorescence in abdomen after exposure to *don juan* males. n = 3 groups of 28 to 32 CS virgin females per time point. **(E-F)** Image captured through the eyepiece of fluorescent microscope demonstrating background signal **(E)** and GFP-positive signal indicating successful mating with *don juan*-GFP male **(F)**. White arrows indicate GFP-positive sperm fluorescence within the abdomen of female flies. **(G)** Average percentage of CS female flies present on sucrose-infused filter paper during a simulated training cycle of 1 min ON and 1 min OFF, repeated 5 total times. Ordinary one-way ANOVA with Dunnett’s multiple comparisons versus the color green. n = approximately 50 flies per group, 5 groups per condition. * = p < 0.05, ** = p < 0.01, and **** = p < 0.0001. **(H)** Average percentage of CS male flies present on sucrose-infused filter paper during a simulated training cycle of 1 min ON and 1 min OFF, repeated 5 total times. Ordinary one-way ANOVA with Dunnett’s multiple comparisons versus the color green. n = approximately 50 flies per group, 5 groups per condition. ns = not significant, *** = p < 0.001, and **** = p < 0.0001. **(I)** Starting (pre-training) preference index for green (PIG) values for CS female and male flies when exposed to green-violet quadrants. Individual circles/squares represent one group of female/male flies. Unpaired t test of starting PIG for females versus males. n = 15 groups of approximately 50 flies per condition. ns = not significant. **(J)** Average (AVG), standard deviation (STDEV), and range for data presented in I. n = approximately 50 flies per group and 15 groups per condition.

In all cases, CS females demonstrated significantly enhanced starvation survival compared to males, which generates disparate levels of motivation for appetitive learning. While the maturation and starvation of flies within same-sex groups produced the most similar responses to starvation, populations consisting of exclusively the same sex do not represent a natural state for flies, which could lead to developmental aberrations and subsequent abnormal behaviors, especially when social behaviors are examined. Non-natural rearing conditions, specifically social isolation, have been shown to affect sleep (Li et al., 2021, Ganguly-Fitzgerald et al., 2006), aggression (Ueda and Kidokoro, 2002), and feeding (Li et al., 2021). The interaction of sleep and memory has been extensively studied in *Drosophila* (Dissel, 2020, Dissel et al., 2015), and indicates that learning can be negatively impacted by impaired sleep (Li et al., 2009, Bushey et al., 2007). As this assay relies on feeding and motivation, in conjunction with learning’s dependence on sleep, unnatural social conditions could negatively impact several aspects of learning in this paradigm.

Based on these data, we suggest using a mixed-sex maturation and starvation in which females are subjected to slightly longer starvation times than males. For our results, this equates to 21 hours of starvation for males and 27 hours of starvation for females, which generates a comparable 80% starvation survival, matching the original protocol’s survival rate. The process of separation of the sexes to accommodate each sex’s distinct starvation time is described in the following optimized assay section. These data also represent the first piece of reasoning for the training and testing of flies in same-sex groups, with additional data supporting this conclusion below.

### Controlling for Sexual Dimorphism in Starvation-induced Dietary Responses

A second sexually-dimorphic factor to consider is that male and female *Drosophila* display differing responses to sucrose. The original protocol used a single sucrose concentration of 1.5 M sucrose on filter paper during the training period. However, when starved, both naïve and mated male flies display increased sensitivity to sucrose and value high carbohydrate food over protein-rich food (Camus et al., 2018). Male flies also exhibit a sucrose-derived tyramine-gated behavioral switch that mediates behavioral decision making when presented with conflicting stimuli (Cheriyamkunnel et al., 2021). When sufficient sucrose has been ingested, tyramine is released, blocking feeding circuits and activating social-behavioral circuits. This same publication shows that at 100 mM sucrose concentrations, starved males switch from feeding to social behaviors after approximately 8 minutes. When compared to the 1.5 M sucrose used in the original appetitive visual STM protocol, it can be assumed that males would switch from feeding to social behaviors much more rapidly, and possibly within the summed 5-minute period of training when sucrose is present. It has also been shown that starvation-induced responses do not reduce social behaviors, even in younger males (Churchill et al., 2019), which suggests that the 4 to 5 day old males used in the experiment will not display quenched social-behavioral motivation upon starvation. Consequently, high sucrose levels can lead to a release of tyramine which may change the fly’s behavioral choice from feeding, and therefore time spent associating a specific color with sucrose, to other social behaviors that obstruct learning. These data suggest that a lower sucrose concentration may be beneficial for male learning by increasing the time spent feeding on sucrose, and thus the association between color and sucrose, while suppressing the switch to other behaviors.

Females also display sexually-dimorphic responses to starvation that are in opposition to that of male flies. Upon starvation, mated females show an increased sensitivity to protein-rich foods, favoring protein sources over carbohydrates, while virgin females do not exhibit such a shift (Camus et al., 2018). Since we are suggesting using a mixed-sex population of flies for maturation and starvation, we examined the mating status of females over 5 days, representing the age of flies at the time of training and testing. Virgin CS female flies were placed into vials with a 1:1 sex ratio with mature (>3 day old) *don juan*-GFP male flies, which carry an endogenous GFP reporter in sperm (Santel et al., 1997). By 72 hours post-introduction, all virgin CS females had been mated in the population, as indicated by a GFP-positive signal within the abdomen of females (Figure 1D-F). Since all females are mated at the time of training and testing, these females should display a higher preference for protein versus sucrose. This suggests that a higher concentration of sucrose may be required for a sufficient response during training, such one comparable to that used in published olfactory memory assays (Tempel et al., 1983, Lyutova et al., 2019, Tonoki et al., 2020), in order to offset the increased preference for proteins.

Based on these data, we suggest that for flies reared in same-sex populations, males and females should be separated into same-sex groups prior to training and testing, and that males should be trained with a lower sucrose concentration (around 50 mM) while females should experience a higher training sucrose concentration (around 2 M). This separation prior to training also mitigates a potential social-behavioral switch of males, as well as resolves the issue of differing starvation times for male and female flies.

### Controlling for the Inherent Color Preference within the Color Combination

In the original protocol, flies were trained and tested on a two-color combination of green and blue. However, recent data have shown that during the daytime, the color green is attractive while blue is exceptionally aversive in *Drosophila* (Lazopulo et al., 2019). Since the original color choice carries such discrepancy, we asked whether there was a more equitable color choice that could be implemented to minimize this inherent color bias. It is possible that if flies are actively avoiding the color blue, they may be failing to sense the presence of sucrose on the filter paper and therefore failing to associate the color with the sucrose reward. To test this hypothesis, we performed a simulated training experiment in which mixed-sex flies were reared for 4 days, starved, and ice anesthetized/separated on the day of the experiment. Males, who were used immediately, experienced 21 hours of starvation and 50 mM sucrose during training, while females, placed back onto starvation medium after separation, experienced 27 hours of starvation and 2 M sucrose during training. Flies were video recorded during the simulated training period to observe the percentage of flies present on the sucrose-infused filter paper over the five consecutive one-minute training sessions.

As predicted by the inherent color bias, sucrose-infused filter paper paired with the color green resulted in the highest percentage of flies present on the filter paper for both males and females (Figure 1G-H). Conversely, sucrose-infused filter paper paired with the color blue results in the lowest percentage of flies present on the filter paper across all colors tested for both males and females. The color violet produced the most similar percentage of flies present on the filter paper compared to green for females (Figure 1G) and the percentages of male flies present on green and violet were indistinguishable (Figure 1H), while red, yellow, and orange exhibited significantly lower percentages of flies present in both sexes when compared to green (Figure 1G-H). Previous publications have shown that low-intensity ultraviolet (UV) light is attractive for *Drosophila* (Heisenberg and Buchner, 1977, Gao et al., 2008, Baik et al., 2018), which may explain why violet, the color closest to UV on the electromagnetic spectrum, is attractive. Additionally, it has been shown that the same histamine chloride channel, *Ort*, is required for preference of both UV and green light (Gao et al., 2008). This finding may not only explain the relative attraction to both colors but may also support the equitability of this color combination, as both use the same channel for color preference. These data suggest that the color combination of green and violet light offers the most unbiased color choice combination, which should generate stronger associations during the training period, leading to more robust learning.

### Controlling for Variation in Pre-training Color Preference with a Pre-training Video

The original protocol recorded a single post-training video to assess the preference index for green (PIG) for each training condition within each test. However, using a single post-training video for comparison assumes that each group of flies has identical, or nearly identical, starting PIG values. Appetitive olfactory memory assays control for this by diluting two odors until there is a 50/50 preference for each odor, i.e., flies are not attracted to or avoiding either odor choice (Tempel et al., 1983, Semelidou et al., 2019). Using this foundation, we asked if the starting PIG values between groups were adequately similar to produce a valid learning result from a single post-training video. To test this, age-matched mixed-sex CS flies were collected, starved, and ice separated into same-sex groups on the morning of the experiment, as previously described. These flies were then placed into a simulated testing protocol, in which a green and violet color quadrant was used with a blank piece of filter paper without sucrose, to determine the PIG in the absence of any training, which we will label as the pre-training PIG.

Both male and female flies displayed similar starting PIG values, still favoring green over violet in the absence of sucrose, reinforcing the attractiveness of the color green (Figure 1I). However, across 15 same-sex groups of both females and males tested, standard deviations of 0.112 and 0.132 and ranges of 0.430 and 0.426, respectively, were observed (Figure 1J). Since the learning indices observed in the original protocol were approximately 0.15, one standard deviation in either direction for a single group is nearly equivalent to the expected learning index. This appreciable variation between groups may conceal learning if only a single post-training video is used to quantify learning.

For example, at two standard deviations, a group of female flies in the green trained condition may display a starting PIG of 0.043 while the violet trained condition has a starting PIG of 0.491. A change of 0.15 in PIG in each condition, which would generate a final learning index of 0.15, results in an ending PIG of 0.193 for the green trained condition and 0.341 for the violet trained condition. A single post-training video of this hypothetical test would show a higher PIG for the violet trained condition and therefore a negative learning index, when in fact, each condition displayed a change in PIG that would be expected with learning. For a graphical demonstration of this situation, see subsequent Figure 2B-E. Overall, this variation in starting PIG values between groups could mask learning and may explain why so many repetitions were required in the original protocol to establish a learning pattern.

**Figure 2:**
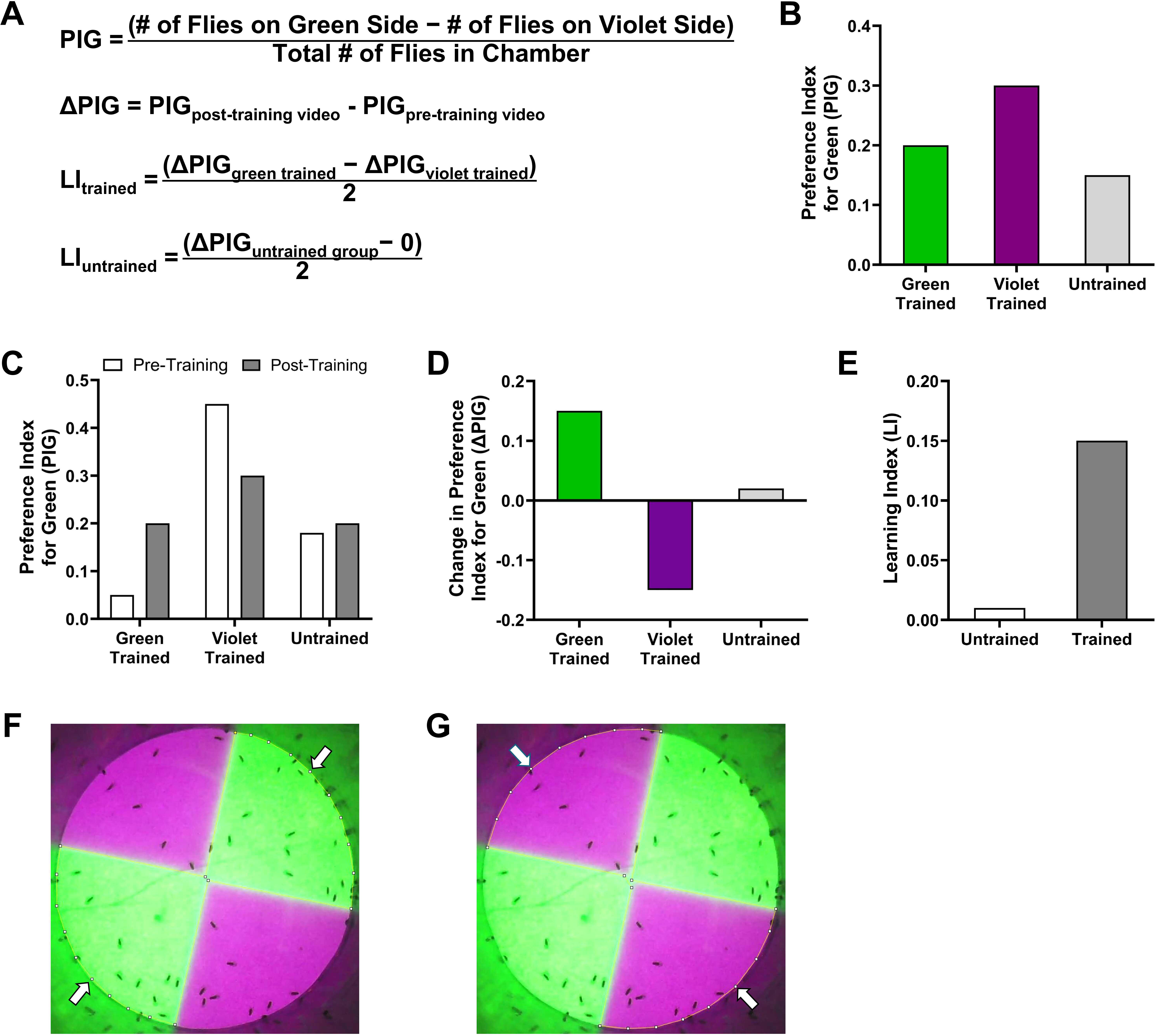
Updated equations and hypothetical experimental results revealing hidden learning. **(A)** Updated equations for calculating PIG, change in PIG, and learning index (LI) using both pre-and post-training videos. **(B)** Simulated result of a single post-training video which does not demonstrate learning if only a post-training video is used. **(C)** Simulated result of the same post-training video results but with addition of pre-training video, demonstrating pre-training differences in color preference. **(D)** Simulated change in PIG values for each condition using the new equations, showing results expected with learning: positive ΔPIG for green and negative ΔPIG for violet, while untrained is near zero. **(E)** Simulated learning index comparing a single post-training video (Post-T Vid Only), which does not show learning, and using both pre- and post-training videos (Pre- and Post-T Vid), which uncovers learning hidden by pre-training color preference differences. **(F)** Green quadrants are selected using ImageJ ROI tool to outline the green area (arrows) prior to initiation of the green-specific ImageJ macro. **(G)** Violet quadrants are selected using ImageJ ROI tool to outline the violet area (arrows). prior to initiation of the violet-specific ImageJ macro.

Based on these data, we suggest implementing a pre-training video to establish a pre-training PIG value as a baseline for each condition, which can then be compared to the post-training PIG value, to more accurately ascertain learning within each condition of flies. The implementation of a pre-training video, however, requires the modification of equations to calculate the learning index.

### Modifications of Equations and Scoring Macros for Quantification

Implementation of the above pre-training video requires the addition and modification of existing equations to quantify learning outcomes. The equation to calculate the PIG value by comparing the numbers of flies on the green and violet sides over the total number of flies, as used in the original protocol only with violet in place of blue, is still employed (Figure 2A, first equation). However, a pre-training video will be recorded for each training condition with a piece of blank filter paper and a pre-training PIG will be calculated for each condition using this equation. After the test, the post-training video will be used to calculate a post-training PIG for each condition using this same equation. These pre- and post-training PIG values will then be used to calculate the change in PIG (ΔPIG) for each condition, using the difference in PIG of the post-training video versus the pre-training video (Figure 2A, second equation). Finally, a learning index can be calculated from ΔPIG values of the green and violet groups (Figure 2A, third equation). The learning index for the untrained group can be calculated by using the ΔPIG for each untrained group and zero, as it is assumed that the untrained groups’ ΔPIG values will be zero in the absence of training (Figure 2A, last equation).

Using the above modifications to equations, a hypothetical experiment was produced to represent the expected results for learning. Similar to when describing the effect of the pre-training variation above, this hypothetical experiment will also show potential masked learning when using only a single post-training video. A single-post training video used to calculate the PIG for each group may indicate that learning did not occur, as the violet trained group has a more positive PIG, i.e., green was favored more than violet (Figure 2B).

However, the implementation of a pre-training video may show that there were learning-derived changes in the PIG values (Figure 2C-D), where a positive ΔPIG was observed in the green trained group and a negative ΔPIG was observed in the violet trained group, signaling that the group learned to associate green and violet with sucrose, respectively. Using these values, a strong learning index is present when using the updated equations but absent in the original protocol where a single post-training video is used (Figure 2E). Overall, these graphs represent the hypothesized patterns expected when learning is present with our optimized protocol.

The original protocol recorded videos at 1 frame per second and calculated the number of flies within each color per second over a 90 second testing period. To assist in scoring, an ImageJ macro was used to count the flies per second, with an error of 2 to 9% over 20 randomly selected quadrants. Following this scheme, we also developed a semi-automated scoring macro within ImageJ using a specific macro for each color to maximize accuracy of scoring, with the only manual step being the outlining of the green or violet quadrants. To score the green trained group, the polygon selection tool was used to outline the green quadrants (Figure 2F). After closing the selection area by connecting the final dot to the starting dot, the green macro was initiated (see Supplementary Material for macro files). The particle count window was saved as a .csv file to be used in Microsoft Excel.

Count data was pasted into an Excel counting template and dead flies were visually counted and subtracted using a built-in equation. This process is then repeated for the violet quadrants (Figure 2G) within each condition. While the threshold values may need to be adapted to different computer monitors and cameras, these macros produced counts with accuracies of approximately 97% for the respective color and were less accurate when either single macro was used for both colors.

Taken together, these data show that the introduction of a pre-training video can identify learning that may be hidden by innate variation in color preference using updated equations and that our semi-automated scoring macros produces highly accurate and expedited scoring. With all the above data taken into account, we can introduce an optimized appetitive visual STM assay that produces significant learning with fewer numbers of flies.

### An Optimized Appetitive Visual Short-term Memory Protocol

Based on the summation of all above data, we successfully implemented and tested female- and male-specific protocols for assessing appetitive visual STM in wild type Canton-S (CS) flies (Figure 3A). Virgin females and naïve males are collected upon eclosion and combined in a 1:1 sex ratio to create age-matched mixed-sex groups. After aging for 4 to 5 days, mixed sex groups are transferred to starvation medium of 1% agar at 12:00pm. The following morning at 8:00am, the groups are ice anesthetized and separated into same-sex groups. Males are trained and tested using 1.5 mL of 50 mM sucrose approximately 1 hour after separation (approximately 21 hours of starvation time) while females are placed back onto starvation medium, to be trained and tested at 3:00pm (approximately 27 hours of starvation time) using 1.5 mL of 2 M sucrose.

**Figure 3:**
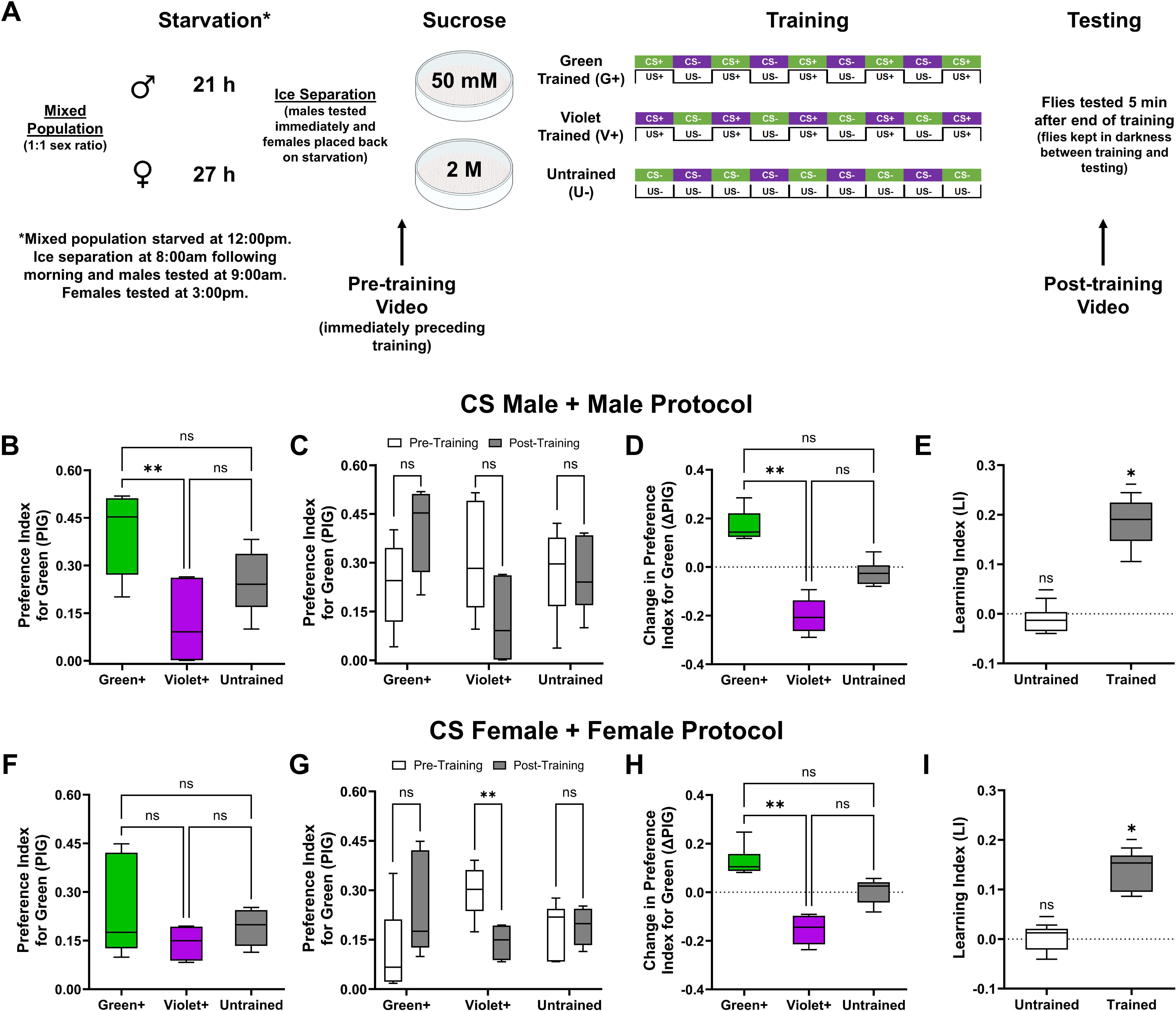
Implementation of the optimized appetitive visual STM protocol. **(A)** Experimental protocol for optimized appetitive visual short-term memory assay with male protocol on top and female on bottom. **(B)** Preference index for green (PIG) only using post-training video data for CS males using the male-specific protocol. Kruskal-Wallis test of multiple comparisons. n = approximately 50 flies per condition for 6 total tests. ns = not significant and ** = p < 0.01. **(C)** Preference index for green (PIG) showing the comparison between the pre- and post-training data for CS males using the male-specific protocol. Mann-Whitney test comparing pre- and post-training PIG values within each condition. n = approximately 50 flies per condition for 6 total tests. ns = not significant. **(D)** Change in preference index for green (ΔPIG) for each condition when comparing pre- and post-training videos for CS males using the male-specific protocol. Kruskal-Wallis test of multiple comparisons. n = approximately 50 flies per condition for 6 total tests. ns = not significant and ** = p < 0.01. **(E)** Learning index (LI) of green and violet (trained) and untrained conditions for CS males using the male-specific protocol. One sample Wilcoxon signed rank test for H_0_: LI = 0. n = approximately 50 flies per condition for 6 total tests. ns = not significant and * = p < 0.05. **(F)** Preference index for green (PIG) only using post-training video data for CS females using the female-specific protocol. Kruskal-Wallis test of multiple comparisons. n = approximately 50 flies per condition for 6 total tests. ns = not significant. **(G)** Preference index for green (PIG) showing the comparison between the pre- and post-training data for CS females using the female-specific protocol. Mann-Whitney test comparing pre- and post-training PIG values within each condition. n = approximately 50 flies per condition for 6 total tests. ns = not significant and ** = p < 0.01. **(H)** Change in preference index for green (ΔPIG) for each condition when comparing pre- and post-training videos for CS females using the female-specific protocol. Kruskal-Wallis test of multiple comparisons. n = approximately 50 flies per condition for 6 total tests. ns = not significant and ** = p < 0.01. **(I)** Learning index (LI) of green and violet (trained) and untrained conditions for CS females using the female-specific protocol. n = approximately 50 flies per condition for 6 total tests. One sample Wilcoxon signed rank test for H_0_: LI = 0. ns = not significant and * = p < 0.05.

Over the course of 6 experiments, CS male flies subjected to the male-specific protocol started to show the expected learning pattern of the original memory protocol (PIG order: green trained > untrained > violet trained), however, the green trained (Green+), violet trained (Violet+), and untrained are not wholly significantly different (Figure 3B).

When including the pre-training PIG data to generate ΔPIG, the anticipated changes in PIG start to emerge, but are still not wholly significant (Figure 3C-D). However, when the ΔPIG values are used to calculate the learning index (LI), significant differences are observed between the untrained and trained groups, where the trained groups show significant learning with a learning index of 0.1906 (Figure 3E). Interestingly, calculations of these experiments using only the single post-training values, as in the original protocol, yields a lower and insignificant learning index of 0.143.

Similarly, CS female flies exposed to the female-specific protocol and tested over 6 experiments produced comparable results, where using a single post-training video to calculate PIG did not show significant differences between any of the conditions (Figure 3F). When using a pre-training video, the expected differences between pre- and post-training PIG values were indicated, although not all groups show significant differences between pre- and post-training PIG values and ΔPIG alone shows separation between the three trained groups, but the differences are still not significant for all groups (Figure 3G-H). However, when using ΔPIG to calculate the learning index, the trained groups show a significant learning index of 0.1534 (Figure 3I), while once again the original protocol using a single post-training video would result in a much lower, insignificant learning index of 0.0489.

To confirm that separate protocols are needed for male and female CS flies, we performed a reciprocal experiment where males were subjected the female-specific protocol (with the exception of starvation times, which was less than 24 hours, as further starvation resulted in significant death), while females were subjected to the male-specific protocol. CS males exposed to the female-specific protocol did not show significant differences in any of the metrics used to calculate learning (Figure 4A-D). Similarly, CS females exposed to the male-specific protocol failed to show significant differences in PIG values, ΔPIG, and learning indices (Figure 4E-H). These data reinforce the need for two separate protocols for male and female CS flies. Taken together, these data show that male and female CS flies exhibit sexually dimorphic behaviors that require sex-specific considerations when performing this appetitive visual STM assay, which can be corrected with the protocol established in this work.

**Figure 4:**
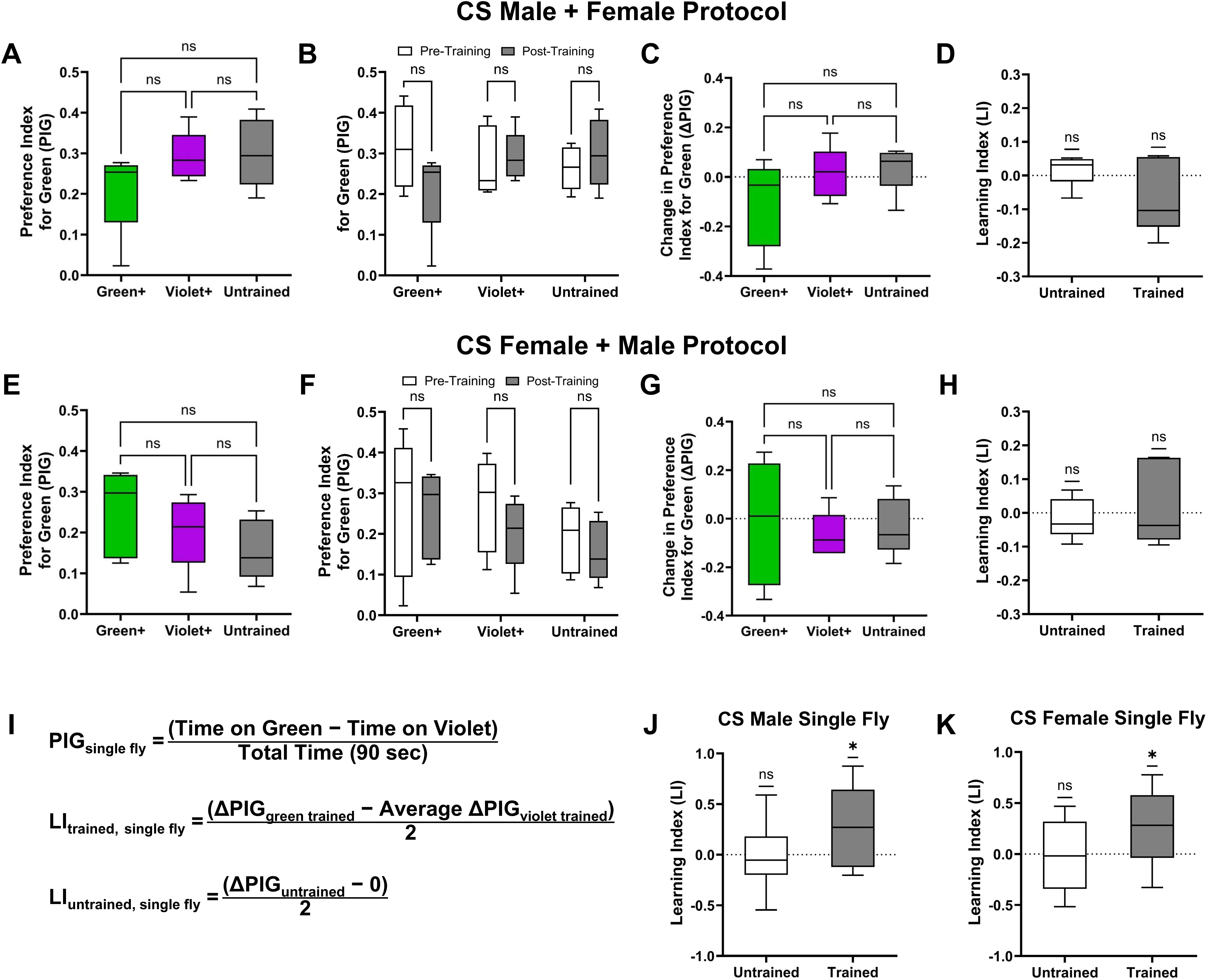
Confirmation of sexually-dimorphic protocol use and single fly validation. **(A)** Preference index for green (PIG) only using post-training video data for CS males using the female-specific protocol. Kruskal-Wallis test of multiple comparisons. n = approximately 50 flies per condition for 6 total tests. ns = not significant. **(B)** Preference index for green (PIG) showing the comparison between the pre- and post-training data for CS males using the female-specific protocol. Mann-Whitney test comparing pre- and post-training PIG values within each condition. n = approximately 50 flies per condition for 5 total tests. ns = not significant. **(C)** Change in preference index for green (ΔPIG) for each condition when comparing pre- and post-training videos for CS males using the female-specific protocol. Kruskal-Wallis test of multiple comparisons. n = approximately 50 flies per condition for 5 total tests. ns = not significant. **(D)** Learning index (LI) of green and violet (trained) and untrained conditions for CS males using the female-specific protocol. One sample Wilcoxon signed rank test for H_0_: LI = 0. n = approximately 50 flies per condition for 5 separate tests. ns = not significant. **(E)** Preference index for green (PIG) only using post-training video data for CS females using the male-specific protocol. Kruskal-Wallis test of multiple comparisons. n = approximately 50 flies per condition for 5 total tests. ns = not significant. **(F)** Preference index for green (PIG) showing the comparison between the pre- and post-training data for CS females using the male-specific protocol. Mann-Whitney test comparing pre- and post-training PIG values within each condition. n = approximately 50 flies per condition for 5 total tests. ns = not significant. **(G)** Change in preference index for green (ΔPIG) for each condition when comparing pre- and post-training videos for CS females using the male-specific protocol. Kruskal-Wallis test of multiple comparisons. n = approximately 50 flies per condition for 5 total tests. ns = not significant. **(H)** Learning index (LI) of green and violet (trained) and untrained conditions for CS females using the male-specific protocol. One sample Wilcoxon signed rank test for H_0_: LI = 0. n = approximately 50 flies per condition for 5 separate tests. ns = not significant. **(I)** Equations for calculating the learning index (LI) of single green and violet trained and test flies (top) and single untrained flies (bottom). **(J)** Learning index for CS males trained and tested individually using the male-specific optimized protocol. One sample Wilcoxon signed rank test for H_0_: LI = 0. n = 17 males per condition. ns = not significant and * = p < 0.05. **(K)** Learning index for CS females trained and tested individually using the female-specific protocol. One sample Wilcoxon signed rank test for H_0_: LI = 0. n = 19 females per condition. ns = not significant and * = p < 0.05.

### *En masse* versus Single Fly Training and Testing

While 6 experiments were needed above to reach significant levels of learning, these still required approximately 900 males and 900 females (150 flies per experiment with approximately 50 flies in each training condition). This number of flies, while substantially lower than the original protocol, is still a considerable number of flies to obtain, especially with respect to crosses involving genetic balancers or in genetic screens. To determine if this optimized protocol generates significant learning for flies trained and tested individually, we reared and starved flies in mixed-sex populations but then subjected single male and female flies to the respective protocol and assessed learning.

Since a single fly was used in a single training condition (unpaired) and the learning index compares green and violet trained flies, modifications to calculating the learning index were required. Following the original protocol, PIG values were calculated by comparing the time spent on each color in seconds over the total time of the test (90 seconds) for both pre- and post-training videos. ΔPIG was calculated as described previously, only for each fly. However, the unpaired nature of the data requires the random pairing of ΔPIG values for calculating the learning index, which can produce dissimilar standard deviations based on the pairings. To mitigate this issue, the learning index for trained flies was calculated using individual ΔPIG values for the green trained flies and the average ΔPIG for violet trained flies (Figure 4I, top equation). The inverse calculation using individual violet trained ΔPIG values and the average green trained ΔPIG will produce an identical learning index with a comparable standard deviation. The learning index for untrained flies was calculated as previously described, this time using individual ΔPIG values for individual untrained flies (Figure 4I, bottom equation).

Both male and female CS flies subjected to single fly training and testing with the sex-specific protocol show significant appetitive visual STM (Figure 4J-K). In each case, the single fly learning indices were higher than those demonstrated using *en masse* conditioning, 0.272 vs. 0.191 for males and 0.255 vs. 0.153 for females, which is consistent with the single fly outcomes in the original publication. While not significantly different from one another at this time, possibly resulting from a small number of *en masse* experiments, further experiments may yield significantly higher learning using the single fly protocol. Importantly, these results were achieved using only 17 male and 19 female CS flies per training condition (less than 60 flies per experiment per sex). These data show that single fly training and testing using optimized sex-specific protocols generates learning at or above the levels of *en masse* conditioning, which greatly reduces the burden of flies required to show significant learning.

## Discussion

*Drosophila melanogaster* has emerged as a powerful tool for the investigation of memory-related processes (Quinn et al., 1974, Schnaitmann et al., 2010, Tempel et al., 1983, Spatz et al., 1974). However, certain paradigms, such as appetitive visual short-term memory (STM), have been less widely used when compared to olfactory and courtship paradigms. Upon implementation of an established appetitive visual STM assay (Schnaitmann et al., 2010), we identified several optimizations based on recent publications that could make the paradigm more efficient, yielding significant learning with fewer repetitions and numbers of flies. We found that in mixed-sex populations, male and female CS flies exhibit significantly different starvation survival (Figure 1B). Since the basis of appetitive memory assays is motivation to find food, this leads to dissimilar motivation levels, and thus learning intensity, between males and females. We also noted that female flies, once mated, will prefer protein-rich foods upon starvation (Camus et al., 2018), while males become more sensitive to sucrose and sucrose-related behavioral changes (Camus et al., 2018, Cheriyamkunnel et al., 2021). In our experiments at the time of testing and training, all females in our mixed-sex populations have been mated (Figure 1D). These data suggested that male and female CS flies require different sucrose concentrations, higher for females and lower for males, to maximize feeding, and thus color association, during the training period. Since mated females do prefer a food source that is rich in protein after starvation, future experiments should investigate amino-acid-infused filter paper, which may generate strong learning in female flies.

We then demonstrated that the color combination of green and blue generates intense bias, and the inherent avoidance of the color blue obstructs learning, with flies actively avoiding the sucrose-infused filter paper paired with blue during training (Figure 1G-H). The color violet produced results the most similar to the color green, and it has been shown that UV-light is attractive for flies (Heisenberg and Buchner, 1977, Gao et al., 2008, Baik et al., 2018), supporting our findings. However, CS flies still showed an inherent preference for green when paired with violet, with significant variations in pre-training PIG values between groups (Figure 1I-J). To account for this inherent variation, we implemented a pre-training video to establish a baseline PIG for each condition prior to training. The introduction of this step also required modification of the existing equations used to calculate learning (Figure 2A).

Using the above data, we generated sex-specific starvation, training, and testing protocols for mixed-sex populations of CS flies (Figure 3A). For both males and females, flies are collected and reared in 1:1 male to female populations prior to separation and starvation. Males experience shorter starvation times, lower sucrose concentrations, and same-sex training and testing. Conversely, females experience longer starvation times, higher sucrose concentrations, and same-sex training and testing. We showed that males display robust learning in the male-specific protocol (Figure 3B-E), while females display robust learning in the female-specific protocol (Figure 3F-I). Interestingly, and in support of the need for sex-specific protocols, males failed to display learning using the female-specific protocol (Figure 4A-D), while females failed to learn using the male-specific protocol (Figure 4E-H). Finally, we also showed that the proper sex-specific protocols generated significant learning in flies that were individually trained and tested (Figure 4J-K), further reducing the number of flies needed to investigate appetitive visual STM.

Our optimized protocols generated significant learning in far fewer repetitions (6 versus 15 or more) when compared to the original protocol that we chose to optimize (Schnaitmann et al., 2010). However, these optimizations do not appear to be universal and should act more as a guide for optimizing this appetitive visual assay in diverse laboratory environments. To explore if the specifications of this paradigm remained valid across different diets, we tested CS flies that were maintained for several generations on a different diet (per 1L: 50 g active yeast, 15 g sucrose, 28 g corn syrup, 33.33 g molasses, and 9 g agar). Compared to our diet used for the optimization process and learning (detailed in Methods), this new diet lacked the presence of cornmeal, was moderately lower in calories and carbohydrates, moderately higher in protein and fiber, and differed in the sources of protein (inactive vs. active yeast) and carbohydrates (presence/amounts of sucrose, corn syrup, and molasses). In doing so, we observed a loss in consistent learning in our optimized appetitive visual STM paradigm (Supplementary Figure 2). It has been shown that diet composition can affect both starvation survival and metabolic processes in flies (Lee et al., 2014, Le Rohellec and Le Bourg, 2009). While some studies show that diet does not influence memory (Ormerod et al., 2017), others have shown that both palatability and nutrient value of food can both influence appetitive memory formation (Burke and Waddell, 2011). Additionally, another study demonstrated that nutrient components of food can alter a fly’s valuation, and therefore choice, of future food opportunities (Das et al., 2016). It has also been shown that the microbiome of the gut, dictated by diet, can also impact memory performance in *Drosophila* (Silva et al., 2021). Taken together, these data indicate that changes in diet alter several factors within the assay itself, including ideal starvation time and sucrose concentrations, that can negatively affect learning.

Our results were found using CS flies, a commonly used stock to represent wild type fruit flies. However, genetic insertions and mutant lines of flies can exhibit differing color preferences, leading to disparity between color choices (Lazopulo et al., 2019), as well as differences in starvation survival (Slade and Staveley, 2016, Lin et al., 1998). If consistent learning is absent in wild type strains, this manuscript can act as a guide and offer recommendations on adapting the protocol, such as starvation time, sucrose concentrations, and color combinations, to control for several possible confounding variables. If learning is evident, but taking many trials to show significant differences, this guide can also help to reduce the number of repetitions needed to show significant learning using pre-training videos and updated equations. Additionally, tests such as starvation survival and the simulated training test with various colors may need to be used to confirm or adapt the assay to meet the needs of various diets and fly genotypes. However, this extra time can be recovered in the significant reduction in numbers of flies and repetitions needed to assess appetitive visual STM with our optimizations.

Sexually dimorphic characteristics of *Drosophila* can impact performance on an appetitive visual STM assay. Factors including starvation survival, starvation-induced dietary changes, and variation of color preferences within the same genotype of flies must be controlled to accurately assess learning. By implementing the changes and protocols detailed above, significant learning can be achieved with a greatly reduced number of flies, especially with the robust learning observed in single fly tests.

## Author Contributions

Conceptualization: S.D., B.L.H, J.A.M

Resources: S.D.

Investigation and Formal Analysis: B.L.H.

Visualization: B.L.H., J.A.M.

Methodology: B.L.H, J.A.M.

Writing – Original Draft: B.L.H, J.A.M, S.D.

Funding Acquisition: S.D.

## Supporting information

Green Macro

Violet Macro

