## Supplementary figures and images for "An optimized appetitive visual short-term memory paradigm in *Drosophila*"

### Supplemental Figure 1

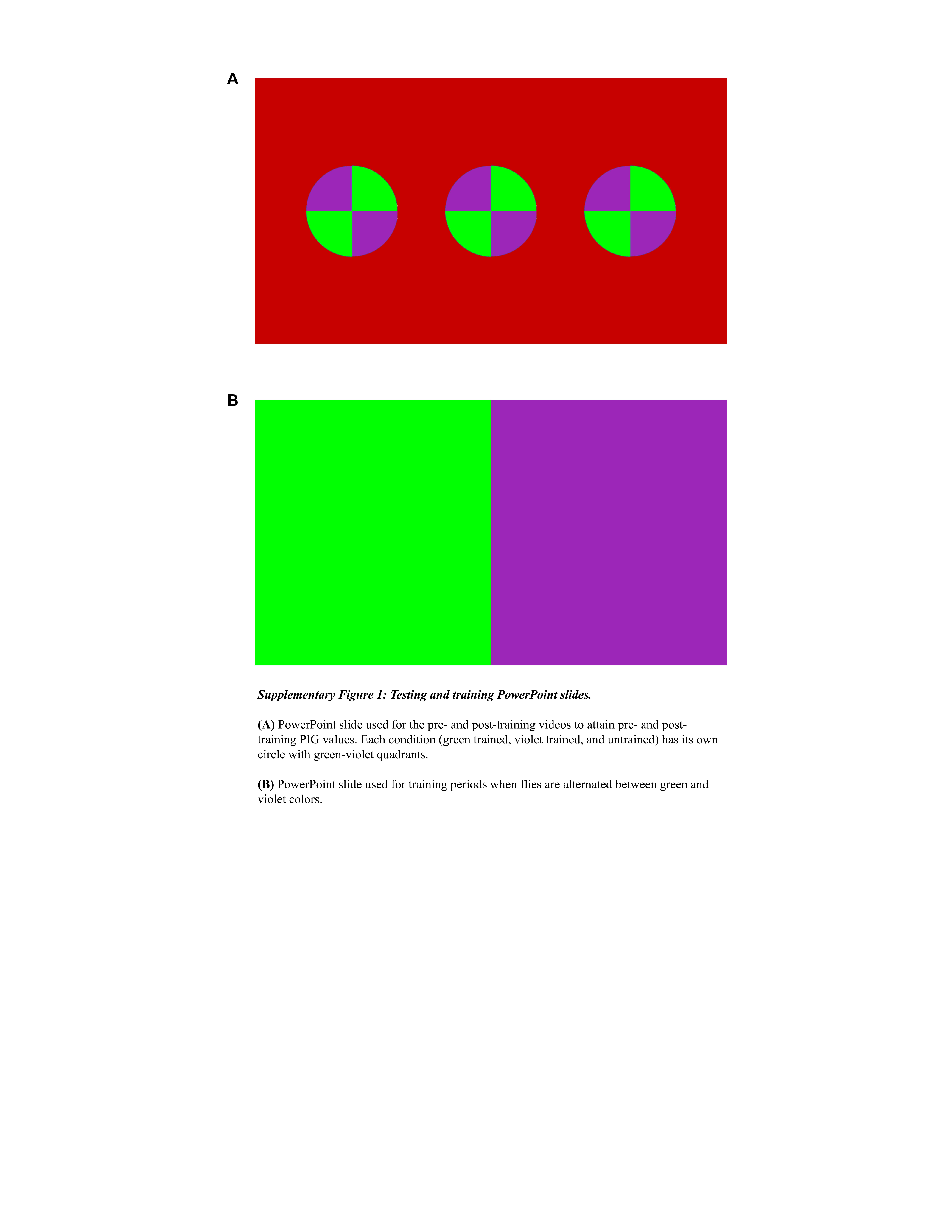

### Supplemental Figure 2

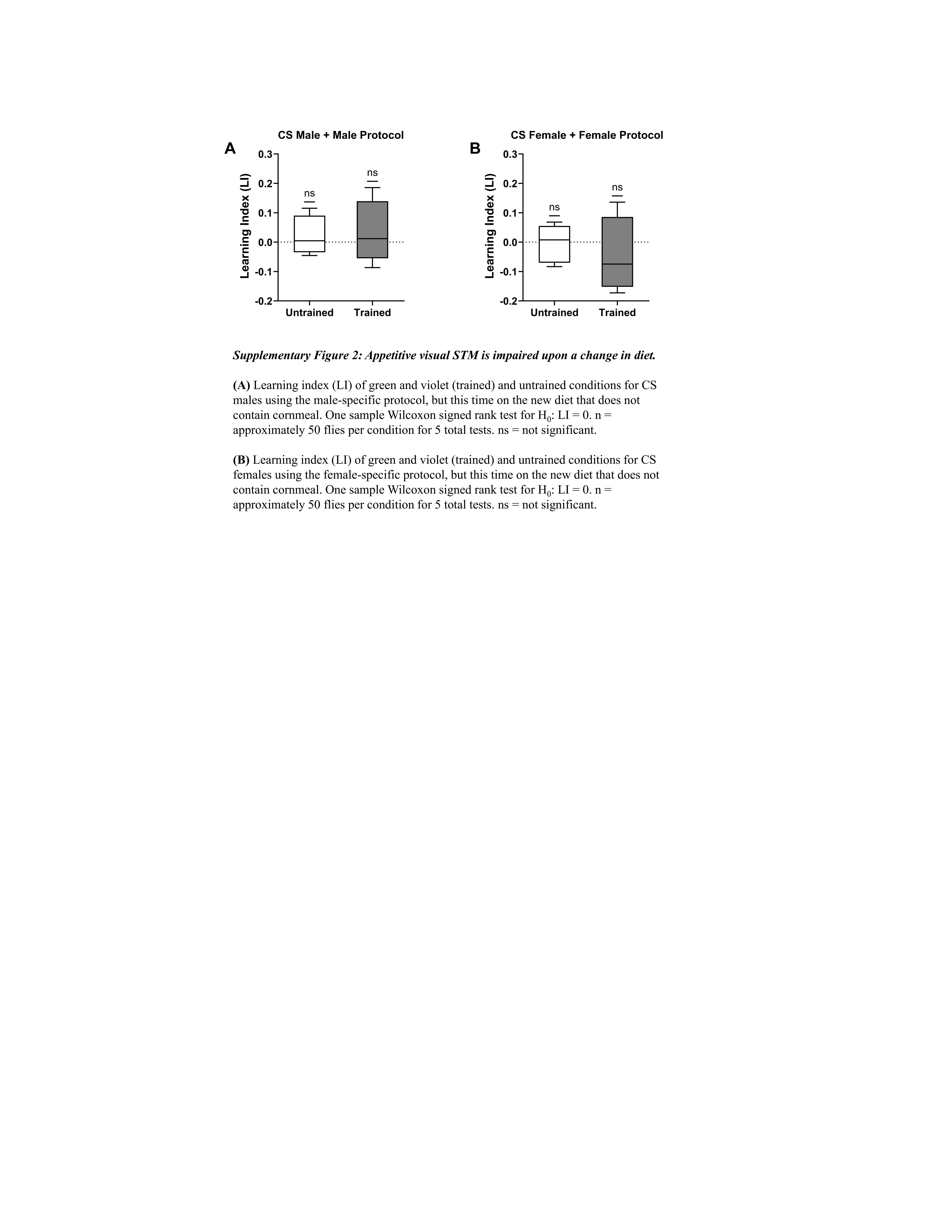
