## Supplementary material for "An optimized appetitive visual short-term memory paradigm in *Drosophila*": Green Macro

run("ROI Manager...");

roiManager("Add");

run("8-bit");

run("Minimum...", "radius=1 stack");

run("Subtract Background...", "rolling=4 light stack");

setAutoThreshold("Default");

//run("Threshold...");

//setThreshold(0, 183);

setOption("BlackBackground", false);

run("Convert to Mask", "method=Default background=Light");

run("Despeckle", "stack");

run("Despeckle", "stack");

run("Despeckle", "stack");

run("Despeckle", "stack");

run("Despeckle", "stack");

run("Watershed", "stack");

roiManager("Select", 0);

run("Analyze Particles...", "size=1-500 circularity=0.25-1.00 show=Outlines summarize stack");
